# Surveillance of zoonotic pathogens in small mammals across a gradient of forest anthropization in Eastern France

**DOI:** 10.1101/2025.06.23.661073

**Authors:** Marie Bouilloud, Maxime Galan, Anaïs Bordes, Caroline Tatard, Philippe Gauthier, Julien Pradel, Guillaume Castel, Hussein Alburkat, Lara Dutra, Tarja Sironen, Clémence Galon, Sara Moutailler, Zorée Djelouadji, Benjamin Roche, Nathalie Charbonnel

## Abstract

The emergence of infectious diseases associated with land-use changes is well-documented. However, zoonotic risks originating from European forests, whether from rural development or urban greening, remain underexplored. To assess and mitigate zoonotic hazards in these ecosystems, we analyzed 1,549 individuals from 18 small mammal species sampled along a forest anthropization gradient using both targeted and broad-spectrum serological and molecular methods. We detected nine bacteria and several Apicomplexa that are potentially pathogenic to humans. Zoonotic pathogen richness and community composition varied significantly across host species, sites and sampling periods. Richness was lower in forested urban parks, possibly due to the absence of vectors or intermediate hosts within cities. It was higher in urban adapter species, even within a given forested habitat, emphasizing the important role of specific life-history traits. Pathogen community structure was similarly shaped by forest anthropization and host ecology, with marked differences between urban and rural forested environments and between urban adapter and dweller species within forested urban parks. The (sero-)prevalence of key pathogens (e.g., *Bartonella, Orthopoxvirus, Neoehrlichia mikurensis*, Sarcocystidae) showed spatial, temporal, and host-specific variation. Site-level differences often exceeded those between general habitat types, highlighting the importance of local ecological context. Nevertheless, some patterns reflected the influence of forest anthropization and species urban adaptation strategies for certain zoonotic agents. Forest anthropization had a positive impact on *Bartonella* prevalence, for urban adapter species within parks, emphasizing a potential dilution effect of these pathogens. Besides, higher levels of Orthopoxvirus seroprevalence were associated with adapter species, in protected forests where they might be more abundant. Altogether, these findings underscore the need for integrated and multi-pathogen wildlife monitoring to anticipate and mitigate disease risks at the human– environment–animal interface.

## 1. Introduction

The recent rise in the number and frequency of zoonotic diseases (re-)emerging worldwide is still concerning (1). It advocates for the implementation of prevention strategies to mitigate these risks (2). While the causes of this rise are complex and multifaceted, land use changes such as deforestation and urbanization seem to play a critical role. They significantly impact environments and host communities, which may result in new favorable conditions for inter-species transmission of pathogens and human exposure to zoonotic agents (3–5).

Forests are biodiversity hotspots and important interfaces between wildlife, domestic animals, and humans, creating opportunities for pathogen transmission (6,7). In Europe, these interactions are amplified by the growing frequency of recreational activities near residential areas (8), but also by ecological pressure that lead to habitat disruption and fragmentation, which displaces wildlife and forces closer contact with humans and domestic animals (5,9).

Despite these growing risks of zoonotic pathogen transmission in European forests, wildlife surveillance remains limited, often focusing on a single known species and pathogen , mainly in response to outbreaks (10). Therefore, it is crucial to implement proactive monitoring of wildlife and zoonotic pathogens to anticipate potential emerging and re-emerging zoonotic diseases, and adapt existing prevention policies to better inform, raise awareness and prepare residents, professionals, and recreational users (11). Besides, the “One Health” approach highlights that the sustainable management of forest ecosystems could offer an effective solution to zoonotic risks, as emphasized in the European Union’s Forest Strategy. However, there remains a significant gap in our understanding of how protecting and restoring forest biodiversity, along with reforesting urban areas, could contribute to mitigating zoonotic risks. This study focuses on the monitoring of multiple pathogens within small mammal communities across forested areas along a gradient of anthropization, covering urban and rural areas (12). As reservoirs of pathogens and bridge species between wildlife and humans (13–15), small mammals constitute a relevant model for pathogen surveillance. Some populations or species may sometimes be declining due to environmental pressures (16,17), yet at other times be thriving in anthropized forest environments (18,19). This positions them as effective sentinels of zoonotic risks. We propose that monitoring small mammals could inform local public health and conservation policies, particularly by identifying species serving as broad-spectrum reservoirs and those inhabiting areas where pathogen transmission to humans or domestic animals is most likely.

First, our objective was to provide a comprehensive description of small mammal communities and their associated zoonotic pathogens in forested areas of Eastern France, encompassing different levels of forest anthropization. We anticipated that analyzing a broad range of small mammal species across diverse forested environments, using multiple pathogen detection methods, would reveal that the zoonotic danger, typically inferred from human cases or existing wildlife surveys focused on particular diseases, has been significantly underestimated.

Second, we analyzed the biotic and abiotic factors that influence the diversity and composition of zoonotic pathogen communities associated with small mammals in this context of forest anthropization. Host species, in combination with sampling sites, are expected to be strong drivers of zoonotic danger as i) small mammal species exhibit varying degrees of competence, *i.e.* capacity to maintain and transmit pathogens (13); ii) environments influence the structure and composition of both host and pathogen communities. More specifically, small mammals’ ecological niche or niche breadth, as well as habitat anthropization, could impact zoonotic hazard, as evidenced by recent meta-analyses. Albery et al. (20) observed that urban-adapted mammal species are associated with a higher number of zoonotic parasites, although this pattern is largely driven by research bias rather than inherent ecological factors. Gibb et al. (4) showed that anthropization and urbanization lead to a significantly higher proportion of known zoonotic hosts ; however their rodent dataset lacked representation from urban environments. We therefore used our dataset to assess the impact of small mammal species’ urban adaptation and the degree of forest anthropization on zoonotic hazard.

Last, we discussed the implications of these results for wildlife surveillance, the need to reconsider prevention policies and guidance on zoonoses risks, all of which are essential for making scientifically-informed decisions to support future forest management and urban greening efforts.

## 2. Materials and methods

All statistical analyses were performed using R v4.4.0 (21).

### 2.1. Small mammal sampling

Small mammal sampling was conducted bi-annually between spring 2020 and spring 2022, in four to six forested sites in Eastern France, encompassing three habitat types: protected forests, managed forests and urban forested parks (Figure 1A). Sampling protocols and study sites are described in detail in Pradel et al. (12). Spatial analyses were conducted to validate the anthropization gradient across study sites, thereby conforming the robustness of the sampling design (Figure S1).

**Figure 1.**
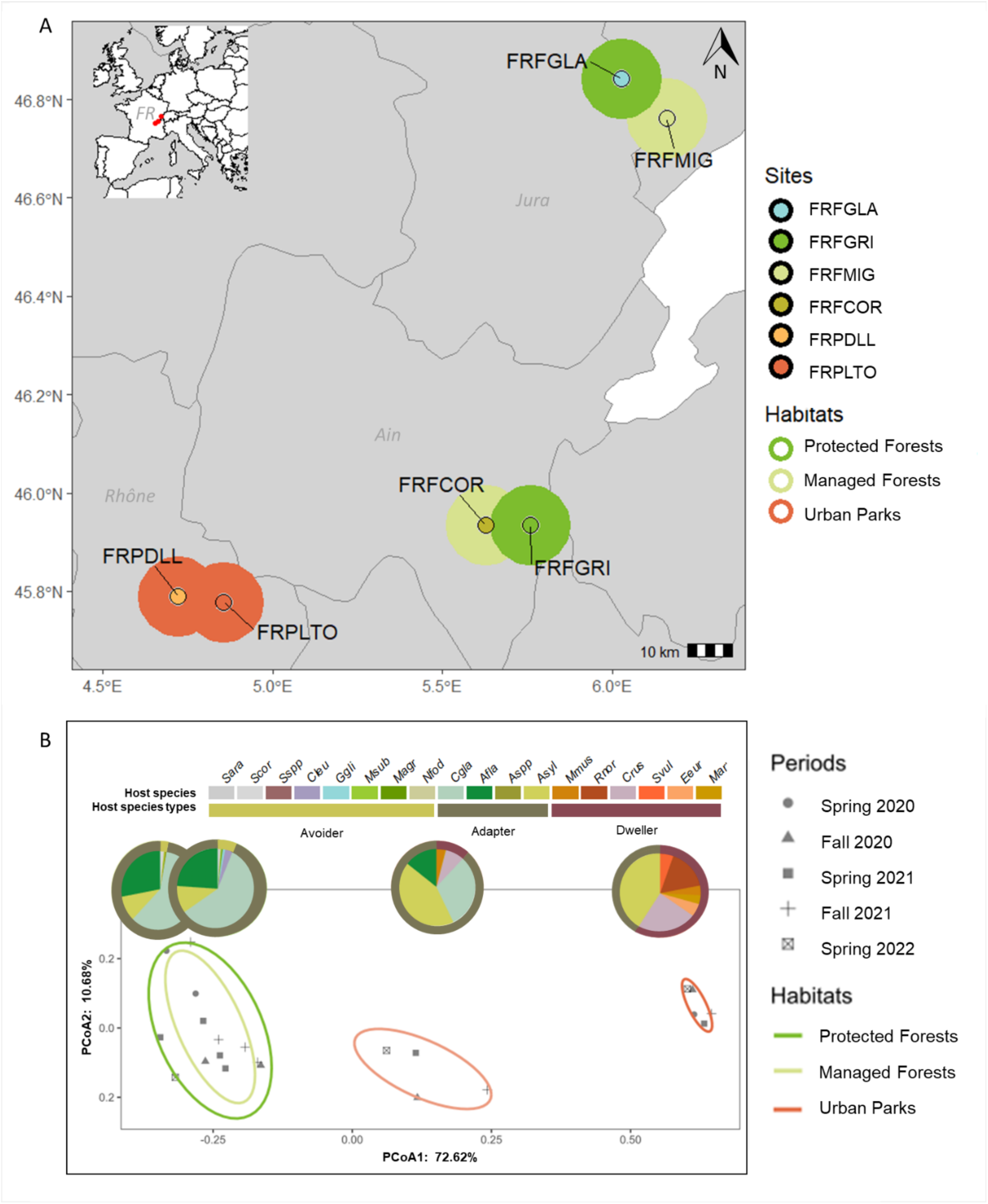
Description of study sites. A. Sampling plan. The inset map shows the study region within Europe, with study sites marked in red. The main map provides a zoomed view of this region in France, displaying the six study sites: FRPLTO: Lyon, Park Tête d’Or (Rhône); FRPDLL: Marcy l’Étoile, Domaine Lacroix Laval (Rhône); FRFCOR: Cormaranche-en-Bugey (Ain); FRFGRI: Arvière, La Griffe au Diable (Ain); FRFMIG: Mignovillard (Jura); FRFGLA: Esserval-Tartre, La Glacière (Jura). Each point is surrounded by a thick border reflecting the associated habitat type: dark green for protected forests (FRFGRI & FRFGLA), light green for managed forests (FRFCOR & FRFMIG), and red for urban forested parks (FRPDLL & FRPLTO). B. Principal Coordinate Analysis (PCoA) illustrating differences in small mammal community composition across study sites and time periods. Each point represents a unique combination of site and sampling period. Different shapes indicate the sampling periods, ordered chronologically from spring 2020 to spring 2023. Ellipses represent habitat types, using the same color code as in the map. Within each ellipse, pie charts illustrate the average relative abundance of each small mammal species. (Species codes: *Asyl = Apodemus sylvaticus, Afla = Apodemus flavicollis, Cgla = Clethrionomys (Myodes) glareolus, Crus = Crocidura russula, Cleu = Crocidura leucodon, Mmus = Mus musculus, Rnor = Rattus norvegicus, Ggli = Glis glis, Msub = Microtus subterraneus, Marv = Microtus arvalis, Magr = Microtus agrestis, Nfod = Neomys fodiens, Svul = Sciurus vulgaris, Eeur = Erinaceus europaeus, Sara = Sorex araneus, Scor = Sorex coronatus).* Species were grouped into urban adaptation categories: avoiders (light green), which avoid urban environments; adapters (dark green), which occur across all habitat types; and dwellers (red), which are exclusively found in urban parks (Figure S2).

Animals were euthanized and dissected to collect individual information (weight, length, reproductive features) and organs necessary for further pathogen detection. Small mammal species were determined based on morphological characteristics, and molecular analyses were applied when necessary (12). The composition of small mammal communities was analyzed through a Bray-Curtis dissimilarity matrix, generated with the package *vegan*, and ordered using Principal Coordinates Analysis (PCoA) with the package *stats*.

### 2.2. Zoonotic pathogen detection

Major rodent-borne disease known in the region were investigated through targeted screening using serological and qPCR approaches, while a metabarcoding approach was used to identify potentially pathogenic bacteria with no *a priori* (22).

#### 2.2.1. Serological analyses for viruses

Blood samples collected from heart in PBS were tested for IgG antibodies against orthohantaviruses (Puumala virus (PUUV) and Dobrava virus (DOBV)), orthopoxviruses (OPXV) and mammarenaviruses (LCMV) using indirect fluorescent antibody tests (IFAT; (23,24)). Positive controls included sera from antibody-positive bank voles for DOBV and OPXV, human for PUUV and LCMV (Progen, Heidelberg). The secondary antibody was a goat anti-mouse IgG conjugated with fluorescein (Jackson ImmunoResearch, Pennsylvania, USA) for all assays but PUUV and LCMV, for which anti-human IgG was used.

#### 2.2.2. Detection of pathogenic *Leptospirosa* species

DNA was extracted from the kidney, target organ for leptospires, of small mammal stored in ethanol with Biobasic kit and pathogenic leptospires were detected using quantitative real-time PCR (qPCR) with Taqman probe (25), targeting the 32-kDa lipoprotein-coding gene (LipL32). All DNA extractions were processed in duplicates, with both positive and negative controls. Using LC480 software, individuals with a C_T_ < 40 were considered positive for pathogenic *Leptospira* species.

Some of the positive samples from all sampling sites and periods, with lowest C_T,_ were genotyped to characterize *Leptospira* species and serovars, as described in Garcia-Lopez et al. (25).

#### 2.2.3. 16S rRNA metabarcoding for bacteria screening

Potential pathogenic bacteria from small mammals were detected in the spleen, an organ involved in pathogen filtration, preserved in ethanol, using the enhanced metabarcoding method of Galan et al. . This approach minimized the risk of false positive results by the validation of the data through systematic technical replicates, filtering thresholds and extensive controls at each step. DNA was extracted using a Qiagen kit, followed by PCR amplification and paired-end Illumina MiSeq sequencing of the V4 region of the 16S rRNA gene (27). The sequences were demultiplexed and analyzed using the *FROGS* (Find Rapidly OTU with Galaxy Solution) pipeline (28). The sequences were grouped using *SWARM* (29) with a maximum clustering distance of d=3 and then compared by *BLAST* to the Silva database v138.1 for taxonomic assignment. The resulting abundance tables were filtered to eliminate potential false positives by applying three series of filters described in Galan et al. . Taxa corresponding to zoonotic pathogens were selected based on the literature.

#### 2.2.4. Specific characterization of bacteria detected by the 16S rRNA metabarcoding

We refined taxonomic resolution by conducting targeted molecular analyses on a small number of 16S-positive individuals to achieve species-level identification within specific zoonotic genera.

Three real-time PCRs were applied for *Francisella* species identification when this genus was detected using the 16S approach (N=70). The ISFtu2-qPCR allowed detecting *F. tularensis* (i.e., *F. tularensis* subsp*. tularensis,* subsp*. holarctica,* and *subsp. mediasiatica*), the Tul4-qPCR enabled detecting *F. novicida* and Type B-qPCR enabled detecting *F. tularensis* subsp. *holarctica* (30).

A subset of *Bartonella* positive DNA extractions (N=11) from four rodent species was sequenced, to achieve a taxonomic resolution to the species level. A two-step PCR protocol, following Galan et al. (31) was applied to amplify two genes, gltA (32) and rpoB (33). Sequence reads were processed using the same FROGS pipeline described above.

The BioMark real-time PCR system (Standard Biotools, San Francisco, CA, USA) was applied to 24 small mammal splenic DNA for high-throughput microfluidic real-time PCR for the most common bacterial tick-borne pathogen species known to circulate or recently emerging in Europe (34,35).

Last, we conducted additional bioinformatics analyses on the 16S rRNA metabarcoding data, focusing on the sequences from apicoplasts associated with *Apicomplexa* from the Sarcocystidae family through BLAST searches and phylogenetic tree construction. A similar species-resolving approach was applied to the sequences of the *Mycoplasma* genus to identify those belonging to the *Mycoplasma* group (Figure S3).

### 2.3. Statistical analyses

We used the *phyloseq* package (36) for data filtering. We compiled pathogen data, keeping only presence/absence information of potentially zoonotic pathogen genera for further statistical analyses. Small mammal species with fewer than 10 individuals were excluded because of the lack of statistical power.

Statistical analyses were then conducted on the finalized dataset to assess how ecological factors shape the presence and transmission dynamics of potentially zoonotic pathogens.

First, the number of pathogens per individual was calculated and modeled using a Poisson distribution (*glmmTMB*, (37)) to account for the high proportion of zeros in the data.

Second, the composition of the pathogen community within individual was analyzed using the Jaccard dissimilarity matrix and a Principal Coordinates Analysis (PCoA) followed by a PERMANOVA test (*adonis2, vegan*). Only individuals carrying at least one pathogen could be considered.

Last, pathogen prevalence was calculated per study site and sampling period, and analyzed using a binomial distribution with a generalized linear model (GLM) for each pathogen with a prevalence greater than 10% in the total dataset. To maximize statistical power and reduce structural zeros, only periods with confirmed pathogen presence were analyzed.

For these three analyses, age class, sex, sampling period, study site and host species were included as explanatory factors. Two modeling strategies were applied. In the first (hereafter referred to as model 1), host species and sites were considered, whereas in the second (model 2), host species were grouped by urban adaptation categories (adapters, dwellers and avoiders), and sites by habitat types (protected forest, managed forest, urban park).

Model selection for GLMs was performed based on AICc (*MuMIn*, δAICc < 2), and model fit was evaluated using DHARMa (38). Multicollinearity between explanatory variables was assessed using the variance inflation factor (VIF). Differences between groups were tested using the post-hoc Tukey HSD test (*multcomp*, (39)) for GLMs, or the multiconstrained *BiodiversityR* (40) for PERMANOVA, with adjustments for multiple testing. To verify species type differences within habitats despite an unbalanced dataset, we compared species separately within the rural (managed and protected) and urban forests subsets. All scripts are available on Zenodo repository.

## 3. Results

### 3.1. Small mammal communities

We collected a total of 1,549 small mammals, including 11 rodent species and seven Eulipotyphla species (*Soricomorpha* or *Erinaceomorpha*) (Figure 1A).

The habitat type, that describes forest anthropization (Figure S1), was the main factor influencing small mammal community composition, with PCoA axis 1 explaining 73% of the total variance. By contrast, sampling periods primarily influenced their abundance (Figure 1B). No difference was found between protected and managed forests, but composition varied significantly between rural forests and urban forested parks.

Small mammal species grouped significantly into urban adaptation categories (Figure S2). Urban adapters (*Apodemus sylvaticus, A. flavicollis* and *Clethrionomys glareolus)* were present and dominant across all environments. Urban avoiders (*Crocidura leucodon, Neomys fodiens, Microtus agrestis, M. subterraneus,* and *Glis glis*) were captured only in rural forests, and at relatively low abundance. In contrast, urban dwellers (*Crocidura russula, Microtus arvalis*, and *Erinaceus europaeus*) were trapped exclusively in urban parks.

### 3.2. Potentially zoonotic pathogens detected

A wide array of pathogenic taxa has been detected, some of them being zoonotic (Table 1).

**Table 1:** Pathogens’ classification and characteristics. This table presents the various pathogens, including bacteria, viruses, and protozoa, that have been detected in this study. Taxonomic details are provided (Kingdom, Phylum, Family, Genus, Species). The mode of transmission is specified as direct or vector-borne, with direct transmission encompassing physical contact or environmental exposure. * indicates that other modes of transmission are known for these potentially vector-borne pathogens. The overall prevalence or seroprevalence is calculated from this study. The principal and potential known diseases associated with each pathogen are indicated.

| Zoonotic Pathogens |  |  |  |  |  |  |  |  |  |
| --- | --- | --- | --- | --- | --- | --- | --- | --- | --- |
| Pathogen Detection and Classification |  |  |  |  |  |  | Pathogen Epidemiological Characteristics |  |  |
| Pathogens codes | Kingdom | Phylum | Family | Genus | Species | Detection Methods | Transmission Mode | Overall Prevalence | Associated Diseases |
| Borr | Bacteria | Spirochaetes | Borreliaceae | Borrelia | *B. garinii*, *B. afzelii*, *B. miyamotoi* | 16S metabarcoding, microfluidic qPCR | Vector | NA | Lyme Disease |
| MycoZ | Bacteria | Firmicutes | Mycoplasmataceae | Mycoplasma | *M. penetrans*, *M. ravidpulmonis* | 16S metabarcoding | Direct | 0.015 | Respiratory Infections |
| Fran | Bacteria | Proteobacteria | Francisellaceae | Francisella | *F. tularensis holarctica* | 16S metabarcoding, qPCRs (ISFtu2, Tul4, Type B), microfluidic qPCR | Vector* | 0.054 | Tularemia |
| Neoe | Bacteria | Proteobacteria | Anaplasmataceae | Neoehrlichia | *N. mikurensis* | 16S metabarcoding, microfluidic qPCR | Vector | 0.121 | Tick-borne fever |
| Rick | Bacteria | Proteobacteria | Rickettsiaceae | Rickettsia | *Rickettsia* spp. | 16S metabarcoding, microfluidic qPCR | Vector | 0.006 | Rickettsioses |
| Anap | Bacteria | Proteobacteria | Anaplasmataceae | Anaplasma | *A. phagocytophilum* | 16S metabarcoding, microfluidic qPCR | Vector | 0.025 | Anaplasmosis |
| Orie | Bacteria | Proteobacteria | Rickettsiaceae | Orientia | *O. tsutsugamushi* | 16S metabarcoding | Vector | 0.019 | Scrub typhus |
| Lept | Bacteria | Spirochaetes | Leptospiraceae | Leptospira | *L. kirschneri*, *L. interrogans* | qPCR (lipL32) | Direct | 0.050 | Leptospirosis |
| Lcmv | Virus | Negarnaviricota | Arenaviridae | Mammarenavirus | *lcmv-like* | Serology (IFA) | Direct | NA | Hemorrhagic fever |
| Hanv | Virus | Negarnaviricota | Hantaviridae | Orthohantavirus | *Puumala virus*, *Seoul Virus* | Serology (IFA) | Direct | 0.060 | Hemorrhagic fever, Nephropathy |
| Poxv | Virus | Poxviridae | Poxviridae | Orthopoxvirus | *Cowpox virus like* | Serology (IFA) | Direct | 0.148 | Cowpox |
| Bart | Bacteria | Proteobacteria | Bartonellaceae | Bartonella | *B. grahamii* (zoonotic), *B. taylorii*, *B. doshiae*, *B. gliris*, *B. birtlesii* and an unknown | 16S rRNA metabarcoding, 2-step PCR (gltA,rpoB), microfluidic qPCR | Vector* | 0.452 | Bartonellosis |
| Chla | Bacteria | Chlamydiota | Chlamydiaceae | Chlamydia | *Chlamydia* spp. | 16S metabarcoding | Direct | 0.012 | Chlamydiosis |
| Toxo | Protozoa | Apicomplexa | Sarcocystidae | Toxoplasma | *T. gondii* | 16S metabarcoding | Direct | 0.001 | Toxoplasmosis |
| Eime | Protozoa | Apicomplexa | Eimeriidae | Eimeria | *Eimeria* spp. | 16S metabarcoding | Direct | 0.002 | Coccidiosis |
| Neis | Bacteria | Proteobacteria | Neisseriaceae | Neisseria | *Neisseria* spp. | 16S metabarcoding | Direct | 0.001 | Meningococci |
| Sarc1 | Protozoa | Apicomplexa | Sarcocystidae | Unknown | Unknown | 16S metabarcoding | Direct | 0.106 | Unknown |
| Sarc2 | Protozoa | Apicomplexa | Sarcocystidae | Unknown | Unknown | 16S metabarcoding | Direct | 0.025 | Unknown |

Serological assays revealed antibodies against four virus families. IFAT detected antibodies against *Orthohantavirus,* and molecular sequencing confirmed the presence of Puumala virus in bank voles from Jura region (10)(Castel et al., 2023) and Seoul virus in brown rats from urban parks (41)(Alburkat et al., 2024). Low seroprevalence was noted for PUUV in mice (1.17%) and DOBV in *Apodemus* species (<0.20%) and *C. russula* (1.04%). Up to now, no

Orthohantavirus could be identified from these seropositive individuals with a molecular approach. A high seroprevalence was observed for Orthopoxvirus (13.19%), whereas Mammarenavirus seroprevalence was low (1.14%). However, no testing was conducted on samples collected in autumn 2021 samples.

qPCR analyses identified pathogenic *Leptospira* bacteria, consistent with their broad host-spectrum reported in the literature. Genotyping revealed a high diversity of *Leptospira* species in urban parks (25), as well as *L. kirschneri Grippotyphosa* and *L. interrogans Australis* in bank voles and *A. flavicollis* from forests.

The 16S rRNA metabarcoding approach was applied to 1,284 individuals. It enabled to detect nine bacteria genera and four protists (Apicomplexa) that are potentially pathogenic for humans and animals (Table 1). Some taxa are obligate vector-borne: *Neoehrlichia, Borrelia, Rickettsia, Anaplasma, Orientia*, while others are or may be spread through direct *(Mycoplasma, Bartonella, Francisella),* potentially sexual (*Chlamydia*) or oral (Sarcocystidae, *Eimeria*, *Toxoplasma*) transmission routes.

qPCR assays on *Francisella* confirmed that 70 metabarcoding-positive individuals were also positive for *F. tularensis holarctica*, the causative agent of tularemia, a high-risk disease in Europe. The sequencing of the rpoB and gltA genes enabled to identify *Bartonella grahamii* (zoonotic), *B. taylorii, B. doshiae, B. gliris, B. birtlesii* and an unknown *Bartonella* species.

The high-throughput real-time PCR approach also enabled to identify *Neoehrlichia mikurensis, Anaplasma phagocytophilum, Borrelia garinii, B. afzelii* and *B. miyamotoi*.

Phylogenetic analyses using the 16SV4 rRNA sequences identified zoonotic variants of *Mycoplasma*, including *M. penetrans* and *M. ravipulmonis* and four taxa groups from *the* Sarcocystidae *family*, with some of the OTUs clustering with *Eimeria sp.* and *Toxoplasma gondii* (Figure S3).

### 3.3. Drivers of zoonotic pathogen presence and transmission dynamics

#### 3.3.1. Richness of zoonotic pathogens per individual

Pathogen richness varied significantly across study sites, periods, and small mammal species (Model 1: Zero-inflated Poisson, Wald test: *sites*: χ^2^ = 44.79, *p* = 1.6×10⁻⁸; *periods*: χ^2^ = 33.53, *p* = 9.3×10⁻⁷; *species*: χ^2^ = 159.60, *p* < 2.2×10⁻¹⁶; AIC = 2968, Figure S4). Considering habitat and species categories (Model 2: AIC = 3178), we found a significant influence of forest anthropization (Zero-inflated Poisson, Wald test: χ^2^ = 7.25, *p* = 2.7×10^−3^) and small mammal urban adaptation (Zero-inflated Poisson, Wald test: χ^2^ = 55.70, *p* = 8.0×10⁻³). Pathogen richness per individual was lower in urban parks than in managed forests (*post hoc Tukey HSD*, β. = – 0.17, *z* = –2.6, *p* = 0.02, Figure 2A), but contrasted patterns were detected for the protected forests FRFGRI and FRFGLA (Table S1, Figure S4A). Urban dweller species exhibited significantly fewer pathogen richness than urban adapter or avoider species (*post hoc Tukey HSD*: adapter - dweller, *β* = –0.86, *z* = –7.48, *padj* < 10^−3^; dweller - avoider, β = –0.77, z = – 2.98, padj = 7×10^−3^, Figure 2B, Table S2). Adapter species harbored about two to three times more zoonotic pathogens than other species categories (Figure 2B). These patterns held even when species co-occurred in the same habitat types (Table S2, Figure S5), although this result was significant in urban parks only (Zero-inflated Poisson, Dweller (ref: Adapter): *β* = -0.85, *z* = -7.28, *p* < 10^−3^).

**Figure 2.**
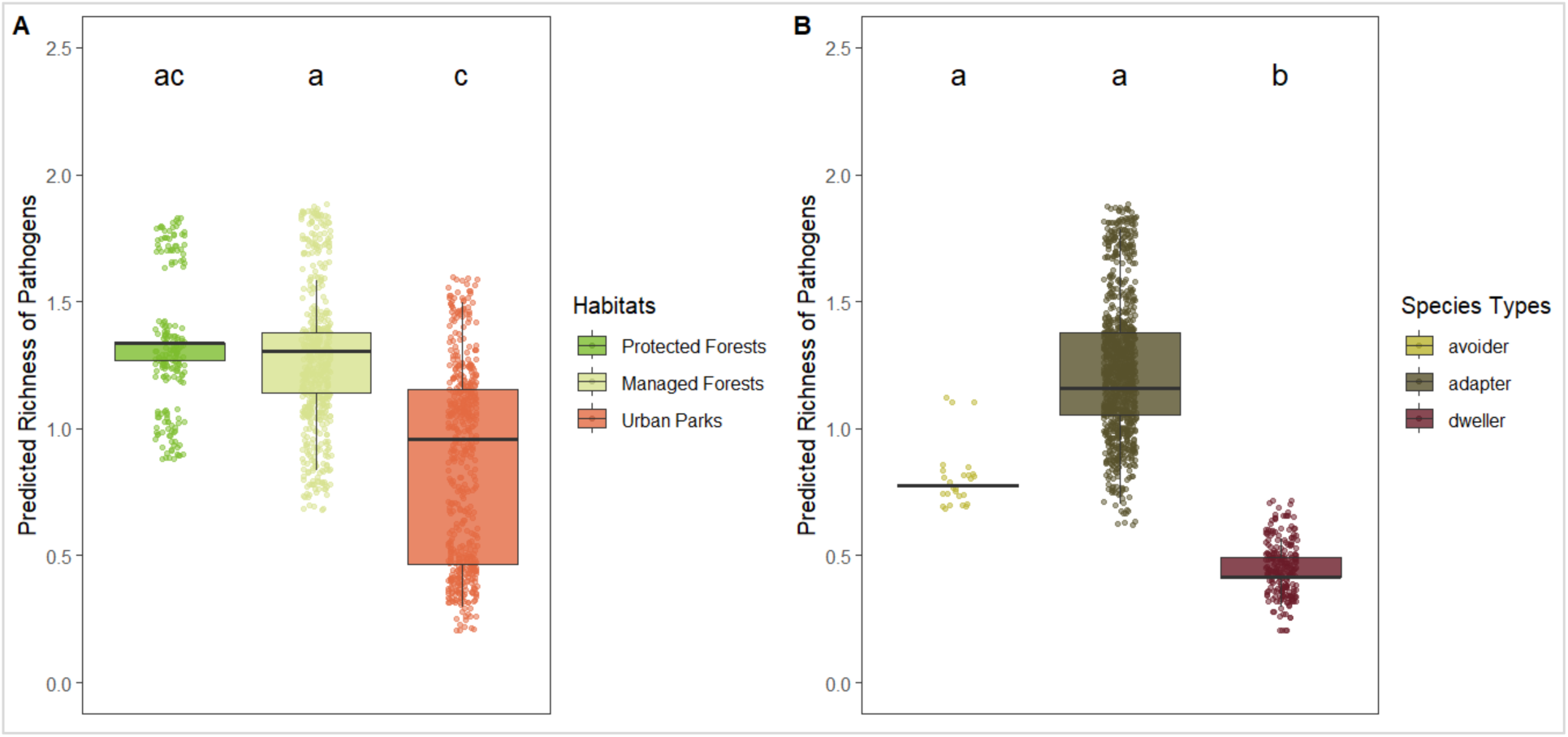
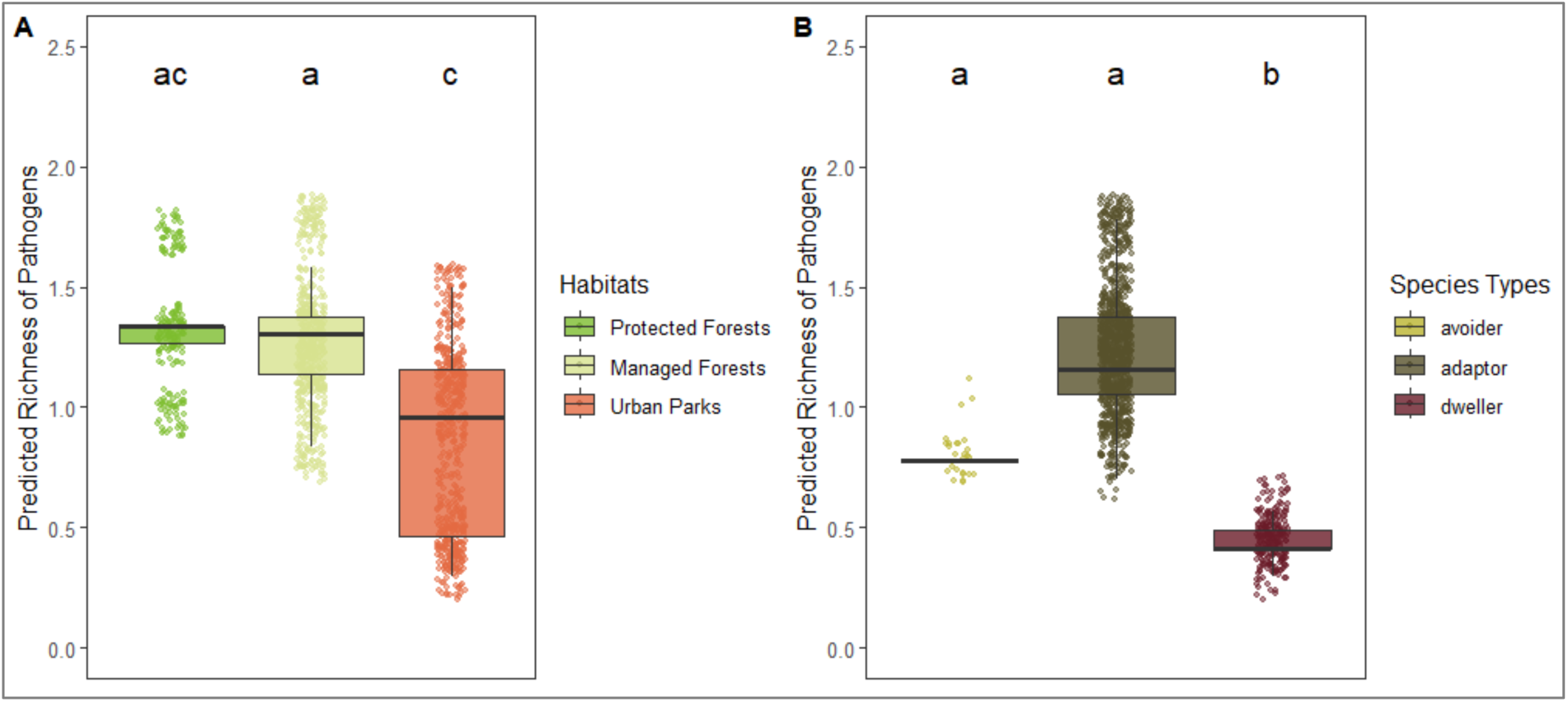
Predicted pathogen richness from *glmmTMB* models **A. across habitat types** (protected forests in light green, managed forests in yellow, urban parks in orange) and **B. by species urban adaptation categories** (avoiders in yellow, adapters in dark green, dwellers in red). Each point represents an individual. Lowercase letters indicate statistically similar or different groups based on post-hoc comparisons. Groups sharing the same letter are not significantly different from each other.

Sampling period significantly influenced pathogen richness per individual, with a significant increase detected from spring 2020 to autumn 2021, and a significant decline in spring 2022 (*post-hoc* tests are detailed in Table S1; Figure S4B).

Among host-related factors, age significantly influenced pathogen richness, with adults harboring more zoonotic taxa than juveniles (Zero-inflated Poisson, *β* = 0.32, *z* = 5.16, *p* < 10^−3^).

#### 3.3.2. Composition of the pathogen community within individuals

PERMANOVA showed that the pathogen community composition within individuals varied significantly according to sampling periods (*R^2^* = 0.04, *p* = 10^−3^), study sites (*R^2^* = 0.07, *p* = 10^−3^), host species (*R^2^* = 0.17, *p* = 10^−3^) and age class (*R^2^* < 0.4%, *p* = 2×10^−3^). Despite their significance, no single factor structured individual groups according to their pathogen composition, reflecting strong inter-individual heterogeneity shaped by host- and context-specific pathogen profiles (Figure S6, Table S3).

Clearer patterns emerged when individuals were grouped based on anthropization-related factors (Figure S7A). Small mammals from managed and protected forests exhibited similar, though weakly differentiated, pathogen communities (SumOfSqs = 3.62, *F*= 1.69, *p* = 10^−3^), whereas individuals from urban parks showed markedly distinct profiles compared to both forest types (urban-managed: SumOfSqs = 6.45, *F*=2.74, *p* = 10^−3^; urban-protected: SumOfSqs = 4.97, *F* = 2.62, *p* = 0.001), as illustrated by the clear separation of habitat types in the PCoA ordination (Figure S7A).

Pathogen communities differed between host urban adaptation categories (PERMANOVA, *R^2^*=0.03, *p*=10^−3^), with the strongest divergence observed between urban adapter and urban dweller species (Figure S7B, Table S4), even within the same habitat (urban parks: adapter– dweller: *R^2^* = 0.12, *p* = 10^−3^; Rural forests (managed and protected) adapter–avoider: *R^2^* = 0.005, *p* = 0.01; Figure 3).

**Figure. 3.**
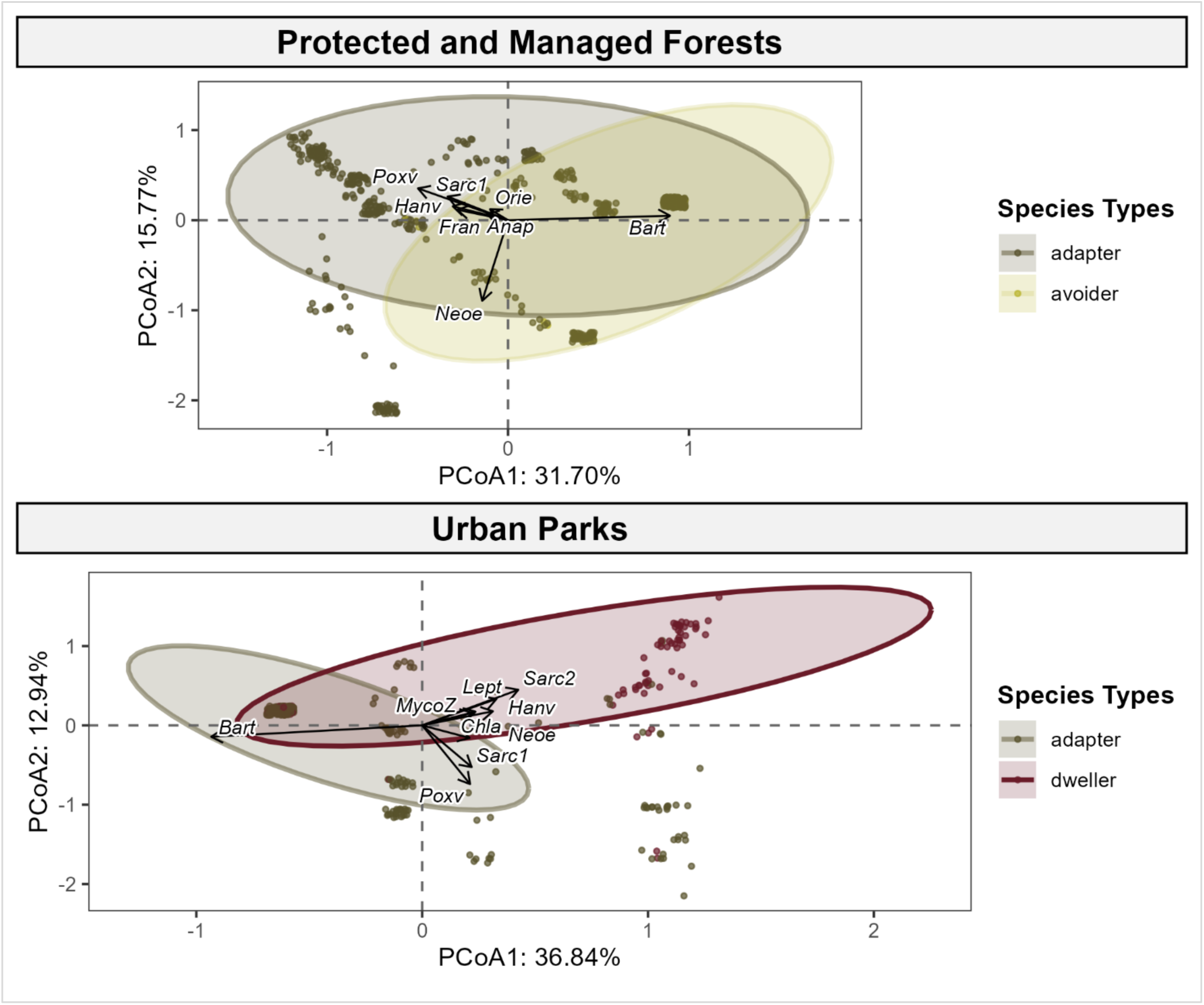
Pathogen composition shown by PCoA based on Jaccard matrix (presence/absence), with points representing individuals (some may overlap). Ellipses (90% threshold) represent species urban adaptation categories in each habitat type (protected and managed forests combined due to similar composition). Environmental vector fitting analysis was performed using *envfit*. Pathogens that were significantly associated with community composition (*p_adj* < 2×10^−3^) are indicated by arrows, showing direction and strength of the signal. Species codes: *Asyl = Apodemus sylvaticus, Afla = Apodemus flavicollis, Cgla = Clethrionomys (Myodes) glareolus, Crus = Crocidura russula, Cleu = Crocidura leucodon, Mmus = Mus musculus, Rnor = Rattus norvegicus, Ggli = Glis glis, Msub = Microtus subterraneus, Marv = Microtus arvalis, Magr = Microtus agrestis, Nfod = Neomys fodiens, Svul = Sciurus vulgaris, Eeur = Erinaceus europaeus, Sara = Sorex araneus, Scor = Sorex coronatus*.

To assess differences in species composition across habitats within an unbalanced dataset, we conducted separate analyses for rural (managed and protected) and urban forest datasets. PCoA biplots highlighted several key pathogens contributing to these variations in pathogen community composition (Figure 3). More specifically, in rural forests, adapter species were associated with higher contributions of *Orthopoxvirus*, *Orthohantavirus,* Sarcocystidae group 1 and *Francisella* sp., and lower contributions of *Orientia* sp. and *Anaplasma* sp., compared to avoiders. Their pathogen communities were strongly shaped by *Bartonella* sp., which exerted a dominant structuring influence. In urban parks, dwellers differed from adapter species by higher occurrences of Sarcocystidae group 2*, Orthohantavirus, Leptospira* sp., and zoonotic *Mycoplasma sp.* Adapter species exhibited higher occurrences of *Bartonella* sp.

#### 3.3.3 Prevalence of zoonotic pathogens

The prevalence of zoonotic pathogens detected in this study strongly varied in space, time and between small mammal species (Figure 4A,B). Further analyses focused on *Bartonella* sp., *Orthopoxvirus*, *Neoehrlichia* sp. and Sarcocystidae group 1, *Francisella turensis*, *Leptospirosa* sp. and *Orthohantavirus* whose (sero)prevalence exceeded 10 % (Table 1).

**Figure 4.**
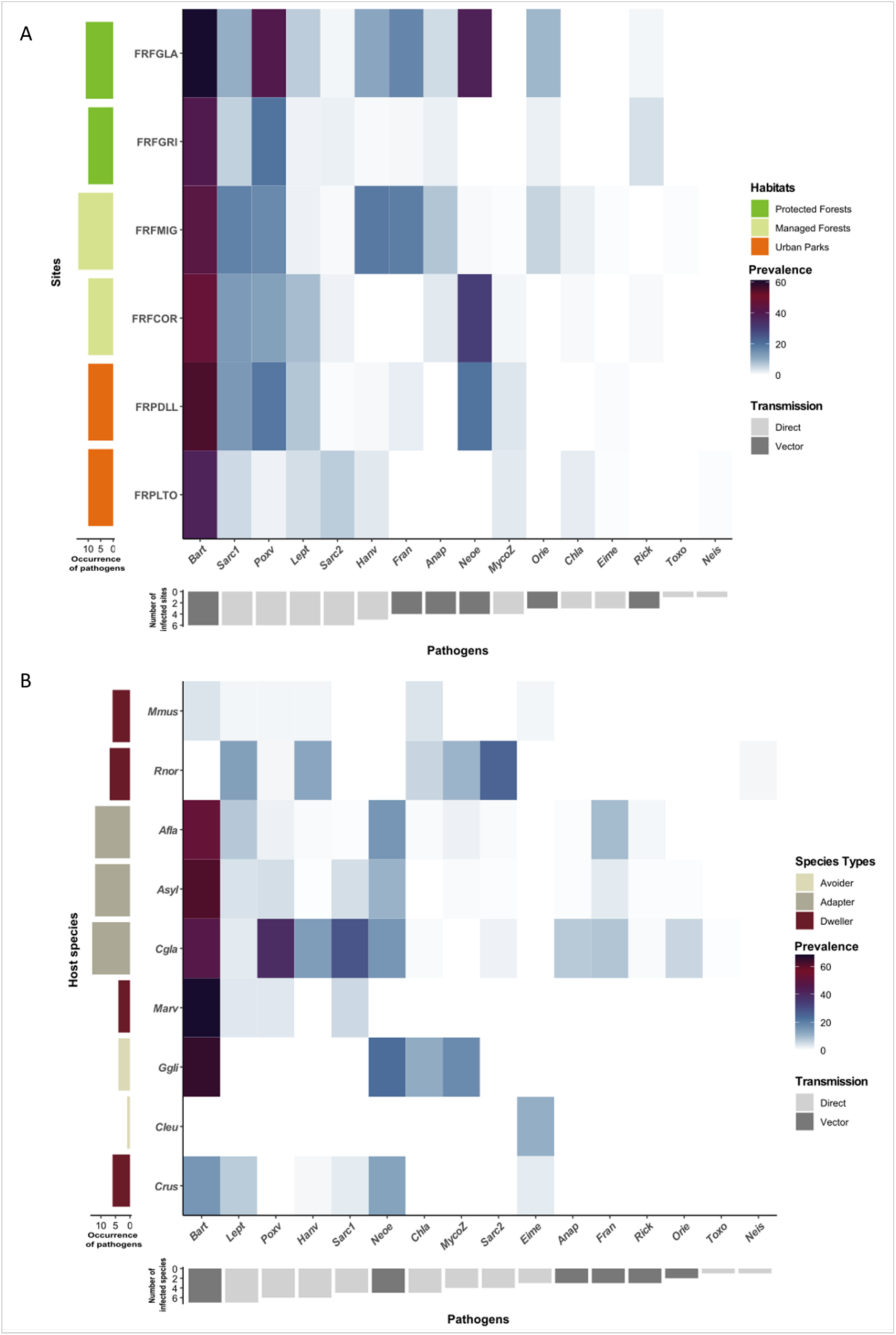
**Heatmap of (sero)prevalence of pathogens: A) by sampling sites** ordered along the anthropization gradient (protected forests: light green; managed forests: yellow; urban parks: orange), and **B) by small mammal species**, grouped by urban adaptation categories (adapters: green-grey; avoiders: light green; urban dwellers: red). Color gradients from light blue to dark purple indicate (sero)prevalence values. Vertical bar charts on the left show the number of zoonotic pathogens detected per site (A) and per host species (B). Inverted horizontal bar charts below indicate the number of sites (A) or host species (B) associated with each pathogen. Pathogens are ordered by prevalence level, respectively per site and host species. Dark grey bars represent obligate vector-borne transmission, light grey bars indicate other potential modes of transmission (via direct, sexual contact, oral routes and/or the environment). Species code: Asyl *= Apodemus sylvaticus,* Afla *= Apodemus flavicollis,* Cgla *= Clethrionomys (syn. Myodes) glareolus,* Crus *= Crocidura russula,* Cleu*= Crocidura leucodon,* Mmus *= Mus musculus,* Rnor *= R norvegicus,* Ggli*=Glis glis,* Marv*=Microtus arvalis*. Site codes : FRPLTO = Parc de la Tête d’Or, Rhône region; FRPDLL = Lacroix Laval, Rhône; FRFCOR = Cormaranche, Ain; FRFMIG = Mignovillard, Jura; FRFGRI = Griffe du Diable, Arvière Ain; FRFGLA = La Glacière, Jura.

*Bartonella* sp. infection, involving a broad host range, varied with host species (ANOVA, model 1, *β* = 268.66, *p* = 2.2 × 10^-16^), with five species being predominantly infected (*Glis glis*, *Microtus arvalis*, *Apodemus sylvaticus*, *A. flavicollis*, *C. glareolus*; prevalence > 40%, Figure 4B), unlike *M. musculus* and *R. norvegicus* (< 5%) (Post-hoc, Table S5). Dweller showed lower prevalence than adapter and avoider species (model 2, post-hoc: *β* = –2.99, *p* < 10^-4^ ; *β* = –3.24, *p* < 10^-4^, respectively). Prevalence also varied across sites (*β* = 46.95, *p* = 5.8x 10^-9^) and sampling periods (*β* = 42.43, *p* = 1.4 × 10^-8^), being higher in urban habitats (urban parks vs. managed forests: *β* = 0.98, *p* < 10^-3^; urban parks vs. protected forests: *β* = 0.66, *p* = 2.7 × 10^-3^, Figure 4A). Peaks occurred in spring 2020 and autumn 2021 (Figure S8; Table S5).

Antibodies against orthopoxviruses showed significant spatio-temporal and species-related variation (ANOVA, model 1: Sites, χ^2^ = 11.70, *p* = 0.04; Species, χ^2^ = 249.05, *p* = 2.2 × 10^-16^; Periods, χ^2^ = 82.839, *p* = 2.2 × 10^-16^), with species urban categories and habitat types significantly explaining prevalence (ANOVA, model 2: Species urban categories: χ^2^ = 37.46, *p* = 7.36 × 10^-9^; Habitats types: χ^2^ = 13.33, *p* = 10^-3^). Antibodies against orthopoxviruses were detected in all small mammal species except *Crocidura* sp. and *G. glis.* Urban adapter species showed higher seroprevalence than urban dwellers (post-hoc test, dweller - adapter: est. = - 2.86, *p* = 2 × 10^-4^). Protected forests had higher prevalence than other habitats (Protected - managed forests: est. = 0.56, *p* = 2.5 × 10^-3^; Urban parks - protected forests: est. = -0.88, *p* = 10^-3^). A peak in seroprevalence occurred in autumn 2021 (Figure S8, post-hoc test, Table S5). The prevalence of *Neoehrlichia* varied significantly between sites (χ^2^ = 257.84, *p* = 2.2 × 10^-16^), with higher rates in FRFGLA and FRFCOR than FRPDLL (Figure 4A ; post-hoc test, Table S5). No significant relationship was found between habitat types (χ^2^ = 5.51, *p* = 0.06). While species alone had no significant impact (not in the best model 1), species categories had a stronger effect (χ^2^ = 13.1, *p* = 10^-3^). Higher prevalence was found in urban avoider and adapter species (post-hoc test, dweller - adapter: *β* = -1.52, *p* = 6 × 10^-3^; dweller - avoider: β = -1.84, *p* = 0.04). Temporal variations were significant (GLM, χ^2^ = 30.42, *p* = 4.03 × 10^-6^), with lower prevalence observed in autumn 2021 (Figure S8, Table S5).

The prevalence of Sarcocystidae 1 varied between species and exhibited significant spatio-temporal variations (ANOVA: Sites, χ^2^ = 17.56, *p* = 4 × 10^-3^; Species, χ^2^ = 204.25, *p* = 2.2 × 10^-16^; Periods, χ^2^ = 32.06, *p* = 1.8 × 10^-6^). It was significantly higher in *C. glareolus* compared to other species (Figure 4B; post-hoc test, Table S5). No patterns were detected based on habitat type or species categories.

## 4. Discussion

### 4.1. A wide diversity of zoonotic hazard

Zoonotic risk associated with small mammals is a global concern, particularly in tropical regions and urban environments where high abundance or diversity of reservoir species, intensified anthropogenic pressures, and ecological or socio-economic conditions favor pathogen transmission to humans (3,4,42). Our analysis highlights a substantial and previously underestimated circulation of rodent-borne pathogens in Eastern France, affecting a diversity of habitats regardless of their level of anthropization.

By combining multiple pathogen detection approaches, we confirmed the presence of known zoonotic agents—such as Puumala virus, *Leptospira* sp., *Borrelia* sp. and *Francisella tularensis* in Eastern France, all of which being already known from wildlife monitoring (10,25,30,43) and human clinical cases (see reports from respective CNR). A more detailed investigation of *Leptospira* in these rural forests revealed the circulation of *L. kirschneri Grippotyphosa* and *L. interrogans Australis*. This provides essential insight for the development of targeted vaccination strategies for professionals like forest workers, who face high occupational exposure risks, especially given that the currently available vaccine only protects against *L. icterohaemorrhagiae*.

Our study also reinforces existing knowledge on the circulation of pathogens that had previously been documented exclusively through wildlife surveillance, including Seoul virus in Lyon (41), *Orthopoxvirus* or *Mammarenavirus* in the Jura region (44).

Importantly, our findings also highlight an overlooked threat with regard to the high diversity of vector borne-bacteria detected in ticks (*Neoehrlichia mikurensis*, *Rickettsia* sp., *Anaplasma phagocytophilum*) and mites (*Orientia tsutsugamushi*). This diversity of vector-borne bacteria underscores the need to expand human diagnostic frameworks beyond *Borrelia* sp. alone, allowing for more accurate recognition of the full spectrum of tick-borne infections that may underlie the wide range of symptoms currently attributed to Lyme disease (45).

While comprehensive, our investigation likely remains incomplete. Important taxonomic groups such as protozoa (e.g. *Giardia* sp., *Babesia* sp.), helminths (e.g. *Echinococcus multilocularis*) and viruses (e.g. flaviviruses, coronaviruses) were not targeted and may represent additional, unrecognized sources of zoonotic hazard. Furthermore, some of the potentially zoonotic taxa we detected, such as *Streptobacillus* sp., *Rickettsia* sp., and Sarcocystidae, warrant further investigation to determine whether they include species of direct relevance to human health.

### 4.2. What shapes zoonotic danger

Our study reveals that small mammal species is a primary factor shaping the diversity and prevalence of zoonotic pathogen communities along the forest anthropization gradient. This finding corroborates previous studies showing that small mammal species exhibit varying levels of reservoir competence (13). It also emphasizes the significance of host-pathogen specificity, which may be driven by phylogenetic relationships, but also by host traits that influence susceptibility, pathogen exposure, and transmission rates. Among them, immunity, behavior or ecological niche have been widely studied (46–48).

Here, we found that urban adapter species, which are generalist species capable of inhabiting a wide range of environments including anthropogenic habitats, harbor zoonotic pathogen communities that differ in composition and exhibit higher richness compared to those of habitat specialists (either urban dwellers or avoiders). Notably, these differences persist even within the same habitat type. This finding aligns with recent meta-analyses suggesting that generalist species tend to allocate more resources in rapid growth and reproduction over immune investment, especially in variable or unpredictable environments, potentially resulting in higher reservoir competence for pathogens (13,49,50).

However, these results have to be considered cautiously as avoider species were represented by a limited number of individuals, due to their low abundance and/or avoidance behavior, while dweller species were difficult to capture and were successfully trapped at a single site only (Lyon urban park). This restricted sampling may introduce bias and limit the representativeness of urban dweller’s zoonotic pathogen communities.

Besides, our analysis underscores that habitat type, here categorized in terms of anthropogenic disturbance, also strongly influences zoonotic pathogen communities. This was mainly observed when comparing urban and rural forests, with cascading effects of forest anthropization on the composition, diversity and interactions of small mammals and zoonotic pathogen communities. These shifts operate through multiple ecological mechanisms, including altered ecological niches, and changes in abiotic conditions and interspecific interactions (47,51).

Although several studies have revealed higher small mammal specific richness in undisturbed habitats compared to anthropogenic ones, due to the exclusion of more specialized, disturbance sensitive species (52), our results rather underscore differences in the composition of these hosts communities. Avoider and dweller species were detected respectively in rural forests and urban parks, whereas urban adapter species were trapped in all types of habitats. These findings align with growing evidence that urban parks may host diverse communities dominated by generalist and synanthropic species, characterized by high ecological plasticity, rapid reproduction rates and behavioral adaptability (53–55). We also showed that urban adapter species thrive in urban parks (e.g. *A. sylvaticus, C. russula*), reaching high densities due to predator release and abundant anthropogenic resources (51,56–58).

These shifts in host community structure, coupled with stressful urban conditions (chemical pollutants and excess artificial light) known to impair immune function and increase host susceptibility, are expected to elevate parasite richness, including zoonotic ones, in urban environments .

Contrary to these expectations, our findings indicate that zoonotic pathogen richness was lower in urban parks compared to rural forests, and not all pathogens exhibited higher prevalence in urban parks. Broader sampling beyond parks might yield different results, but several factors could explain this pattern. First, urban-adapted mammal species are not necessary linked to more zoonotic parasites (see impact of publication bias, (20)), emphasizing the need for more field-based data. Second, the constant availability of anthropogenic food in urban parks may decrease foraging and exposure to environmentally transmitted parasites (60). Third, urban-induced changes in microclimate and vegetation, such as reduced canopy cover and increased forest fragmentation, may limit the diversity or abundance of vectors and intermediate hosts (61). A recent meta-analysis supports this, showing that parasites with complex life cycles were less prevalent in urban dwelling primate and carnivores compared to less anthropized habitats, in line with the urban refuge hypothesis (55). Similarly, our study showed a decline of vector-borne zoonotic pathogens such as *Rickettsia* sp., *Anaplasma* sp. and *Orientia* sp. in urban parks, though *Neoehrlichia mikurensis* was an exception, emphasizing the complex and multifaceted impacts of forest anthropization on zoonotic hazards.

Our results also suggest that specific ecological interactions between hosts and pathogens may play a key role in determining how anthropization influences zoonotic hazard, leading to idiosyncratic patterns. For example, *Bartonella* sp. were highly prevalent across host species and sites, with higher prevalence in urban adapter species, and in urban parks. This pattern may reflect environment-driven changes in host behavior, exposure or susceptibility, though this trend was not observed for other pathogens, suggesting that *Bartonella*’s ecology plays an important role. Factors like *Bartonella* species diversity, coinfections (62), or environment-dependent transmission (63) may be involved. These findings highlight the complexity of host– pathogen dynamics in anthropized landscapes and the need to consider pathogen-specific traits in zoonotic hazard assessments.

Differences between protected and managed rural forests appeared more subtle than expected, potentially due to confounding ecological and climatic characteristics. Shared abiotic features could explain the similarity of host communities’ composition. Both forest types were dominated by urban adapters, with low abundance of avoiders. Moreover, site-specific characteristics may blur distinctions: for example, ’La Glacière’ reserve is small and bordered by managed forests, likely experiencing similar pressures than the close managed forests of Mignovillard. Still, some pathogens showed varying prevalence. *Chlamydia* sp., zoonotic *Mycoplasma* sp., and Puumala virus tended to be less prevalent in protected forests (10), while *Orthopoxvirus* seroprevalence was higher—possibly due to greater biodiversity or older host populations increasing exposure (64).

Lastly, fine-scale spatial and temporal variations in zoonotic pathogen prevalence were observed, with particular taxa being detected only once during this multi-annual survey (e.g. *Francisella tularensis*). Local abiotic conditions, mast seeding, may boost small mammal abundance and in turn, the transmission rate of certain pathogens (e.g. for *Orthohantavirus*, (65,66)). These findings underscore the need for longitudinal and spatially resolved studies to capture transient yet ecologically meaningful shifts in zoonotic hazard, particularly in the context of ongoing environmental changes.

### 4.3. Consequences for zoonotic hazard mitigation

The observed spatial and temporal heterogeneity in zoonotic pathogen richness and prevalence across rural forests and urban parks calls for tailored surveillance strategies and context-specific public health responses. Regardless of habitat types, targeted public awareness is a critical component of effective risk mitigation.

Several zoonotic pathogens such as *Leptospira* sp., Orthohantavirus (SEOV), and *Bartonella* sp., were found at high prevalence in urban wildlife, particularly in adapter species such as *Apodemus sylvaticus* and *Rattus norvegicus*. These species thrive in close proximity to humans, providing opportunities for exposure and potential spillover risk despite lower overall pathogen richness in urban parks compared to rural forests. Urban parks, where high human densities and activity intersects with abundant competent reservoir hosts, therefore represent critical zoonotic interfaces (67). With expanding urban greening initiatives, proactive measures including surveillance and targeted prevention strategies will become increasingly important (68).

Surveillance should focus on sentinel species, particularly synanthropic adapters, that serve as main competent hosts for key zoonotic pathogens (54), and incorporate integrated rodent management strategies. These include improved waste management, biodiversity restoration to promote dilution effects, and physical buffers separating human recreational areas from rodent habitats. Interventions should be concentrated in high-risk areas with frequent human or captive animal contact with infected rodents. Altogether, these proposals should provide local environmental and Public health stakeholders with science-based tools for implementing evidence-driven interventions (69).

In rural forests, a considerable diversity of rodent-borne zoonotic pathogens, in particular vector-borne ones, was detected. Even though human and domestic animal exposure may be lower, the need to mitigate zoonotic risks remain significant. Continuous surveillance, public education and forest management are needed to reduce human-wildlife contact and support ecosystem health. Monitoring should capture seasonal and annual fluctuations in host populations and pathogen circulation, forming an early warning system for emerging threats. Effective surveillance must be large-scale to assess the diversity of host and vector species. This should include non-invasive wildlife survey (e.g., camera traps, environmental DNA), environmental data (e.g., temperature, humidity), and pathogen screening.

From a management perspective, factors like forest fragmentation, habitat connectivity, and land use must be considered in designing interventions to mitigate zoonotic risk (70). Even without strong evidence for the dilution effect here, ecological restoration and biodiversity conservation may still confer indirect protective benefits. These include buffering against the establishment of invasive species and supporting predator populations that naturally regulate rodent densities. These ecological processes can ultimately reduce the abundance of reservoir species and associated zoonotic hazards.

## 5. Conclusion

This study reveals a high diversity of pathogens circulated within small mammal communities across both urban and rural forests in Eastern France. Zoonotic hazard emerges from complex interactions between host species traits and the degree of habitat anthropization. These results highlight the need for developing habitat-specific research and surveillance programs that consider seasonal fluctuations, host ecology, and environmental changes. Mitigation efforts should integrate public awareness, ecological restoration, and targeted management of reservoir species. Ultimately, strong collaborations between researchers, environment managers, clinicians, and public health authorities will be critical to anticipate, detect and reduce zoonotic risks in an increasingly human-altered landscape.

## Ethics Statement

The CBGP laboratory has approval (F-34-169-001) from the Departmental Direction of Population Protection (DDPP, Hérault, France) for the sampling of small mammals and the storage and use of their tissues. All procedures related to small mammals captured in this study complied with the ethical standards of the relevant national and European regulations on the protection of animals used for scientific purposes (Directive 2010/63/EC revising Directive 86/609/EEC, adopted on 22 September 2010). All procedures have undergone validation by the regional ethics committee “Comite d’Ethique pour l’Expérimentation Animale Languedoc Roussillon n°36”in 2020 (ref: 2020-02-v2).

## Acknowledgments

The authors acknowledge the people from ONF (especially Julien Desbois, Thomas Benoit, Nicolas Micoud), the park ‘Lacroix-Laval’ (Julie Dussert), ‘la Tête d’Or’ (Diana Sepulveda, Fabrice Delaveau, Cédric Verjat) and the zoological park of Lyon city (Gwendoline Anfray, Alban Chauvet) for their help in the field. We would also like to thank the GenSeq technical facilities of MEEB (CNRS and University of Montpellier) hosted by ISEM (CNRS, University of Montpellier and IRD) and Audrey Weber (INRAE-AGAP) for the MiSeq sequencing runs, and the Genotoul bioinformatics platform Toulouse Midi-Pyrénées (Bioinfo Genotoul). The authors thank Nora Madani for conducting the genetic analyses used in the characterization of *F. tularensis*.

## Supporting Information

Additional supporting information can be found online in the Supporting Information section.

Supporting Information 1.

Table S1. Richness of pathogen taxa per individual according to species and sites (Model 1) Table S2. Richness of pathogen taxa per individual according to species categories and habitat types (Model 2)

Table S3. PERMANOVA showing differences in community composition (Jaccard distance) according to sites and species (Model 1)

Table S4. PERMANOVA showing differences in community composition (Jaccard distance) according to Habitat types and species categories (Model 2)

Table S5. GLM analyses of pathogen prevalence according to sites and species (Model 1), and habitat types and species categories (Model 2)

Supporting Information 2.

Fig. S1. Principal Component Analysis (PCA) of study sites based on their biogeoclimatic (A) and anthropogenic (B) characteristics.

Fig. S2. Principal Coordinates Analysis (PCoA) based on the distribution of small mammal species across study sites.

Fig. S3. Phylogenetic trees of Sarcocystidae and *Mycoplasma* based on 16S V4 sequences.

Fig. S4. Boxplots of predicted individual pathogen richness from zero-inflated Poisson GLMs, according to: A. sites, ordered along an anthropization gradient; B. periods, C. small mammal species.

Fig. S5. Boxplots of predicted individual pathogen richness from zero-inflated Poisson GLMs, according to the ecological types of small mammals along the anthropization gradient.

Fig. S6. Pathogen community composition (presence of at least one pathogen among the 16 tested) based on Jaccard dissimilarities visualized by PCoA, colored according to site, sampling period, and host species.

Fig. S7. Pathogen community composition (presence of at least one of the 16 pathogens) based on Jaccard dissimilarities.

Fig. S8. Heatmap of pathogen (sero)prevalence across sampling periods. Sampling periods are ordered chronologically.

## Data Availability Statement

16S data from splenic DNA are available at https://zenodo.org/records/12518286

Pathogen characterization data and scripts are available at https://10.5281/zenodo.15671364

