## Supplementary Figures for "Surveillance of zoonotic pathogens in small mammals across a gradient of forest anthropization in Eastern France"

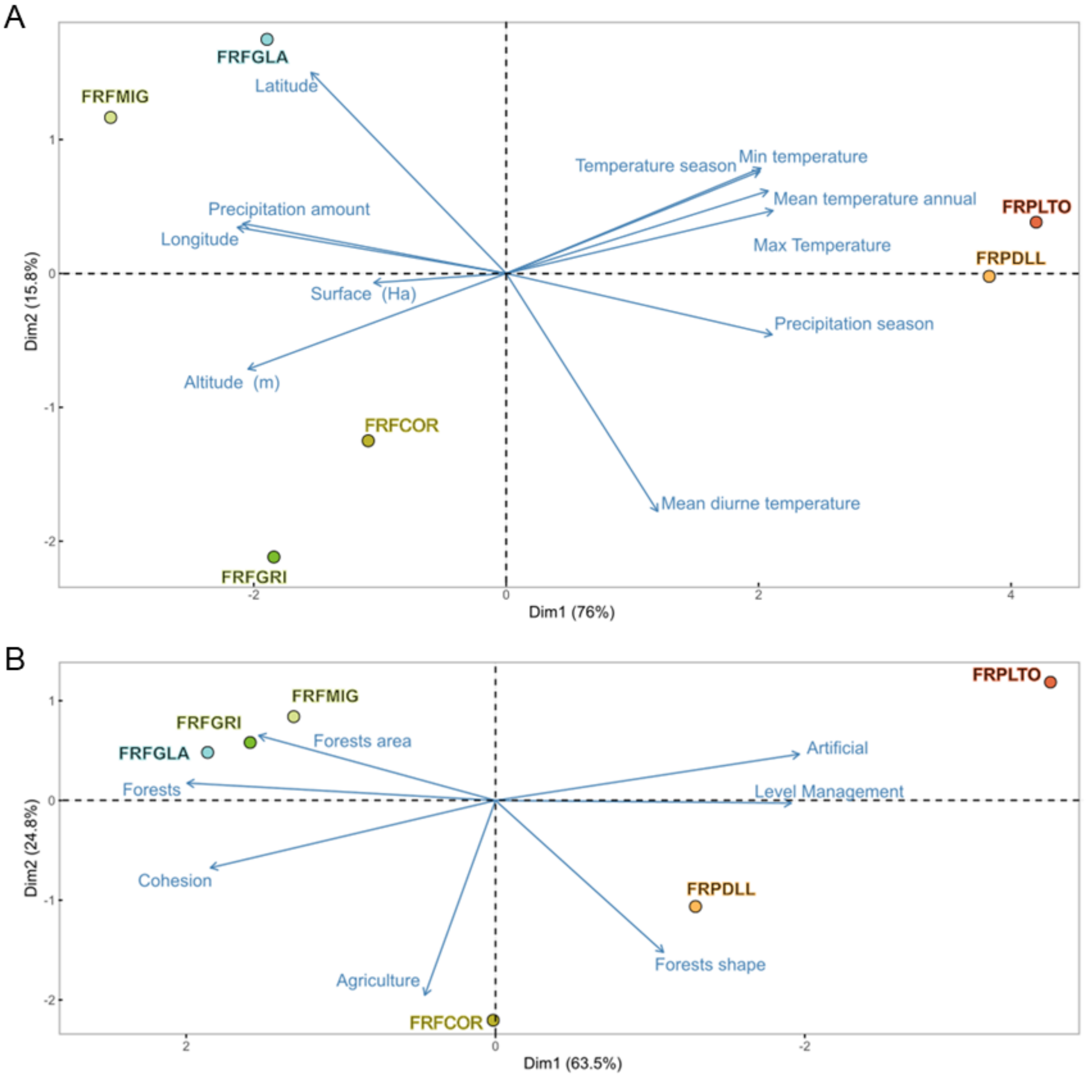


**Supplementary Fig. 1. Principal Component Analysis (PCA) of study sites based on their biogeoclimatic (A) and anthropogenic (B) characteristics.** Analyses were conducted on centered and scaled matrices using the rda function from the vegan package in R (Oksanen et al. 2020). **A. PCA of biogeoclimatic characteristics:** Various factors are derived from GPS coordinates and the Chelsa database (definitions on this site (Oksanen et al. 2020)https://chelsa-climate.org/bioclim/); **B. PCA based on anthropogenic factors.** FRPLTO: Lyon, Parc de la Tête d'Or (Rhône); FRPDLL: Marcy l'étoile, Domaine Lacroix Laval (Rhône); FRFCOR: Cormaranche en Bugey (Ain); FRFGRI: Arvière, La Griffe au diable (Ain); FRFMIG: Mignovillard (Jura); FRFGLA: Esserval-Tartre, La Glacière (Jura). Blue arrows represent the different variables, and the points and annotations vary in color according to the anthropization gradient, with red indicating the most urban areas. The first principal component of the PCA on bio-geoclimatic characteristics explained 76% of the variance, primarily distinguishing sites by longitude, precipitation, and temperature. Study sites from the Rhône region (FRPLTO and FRPLL) had higher temperatures than those from Jura and Ain. The second PCA axis, explaining 15% of variance, revealed variations in latitude and altitude, with study sites from Ain showing higher altitude and lower latitude, leading to warmer daytime temperatures. The first principal component of the PCA on anthropogenic characteristics explained 63% of variance, contrasting managed artificial zones with larger forest areas. Study sites described an anthropization gradient, with urban sites (FRPLTO, FRPDLL) contrasting with rural ones (FRFCOR, FRFMIG). This axis generated a quantifiable anthropization gradient score. The second axis accounted for 29.83% of variance, indicating land management pressures, contrasting agricultural and fragmented forest areas (FRFCOR) with denser forests (FRFMIG).

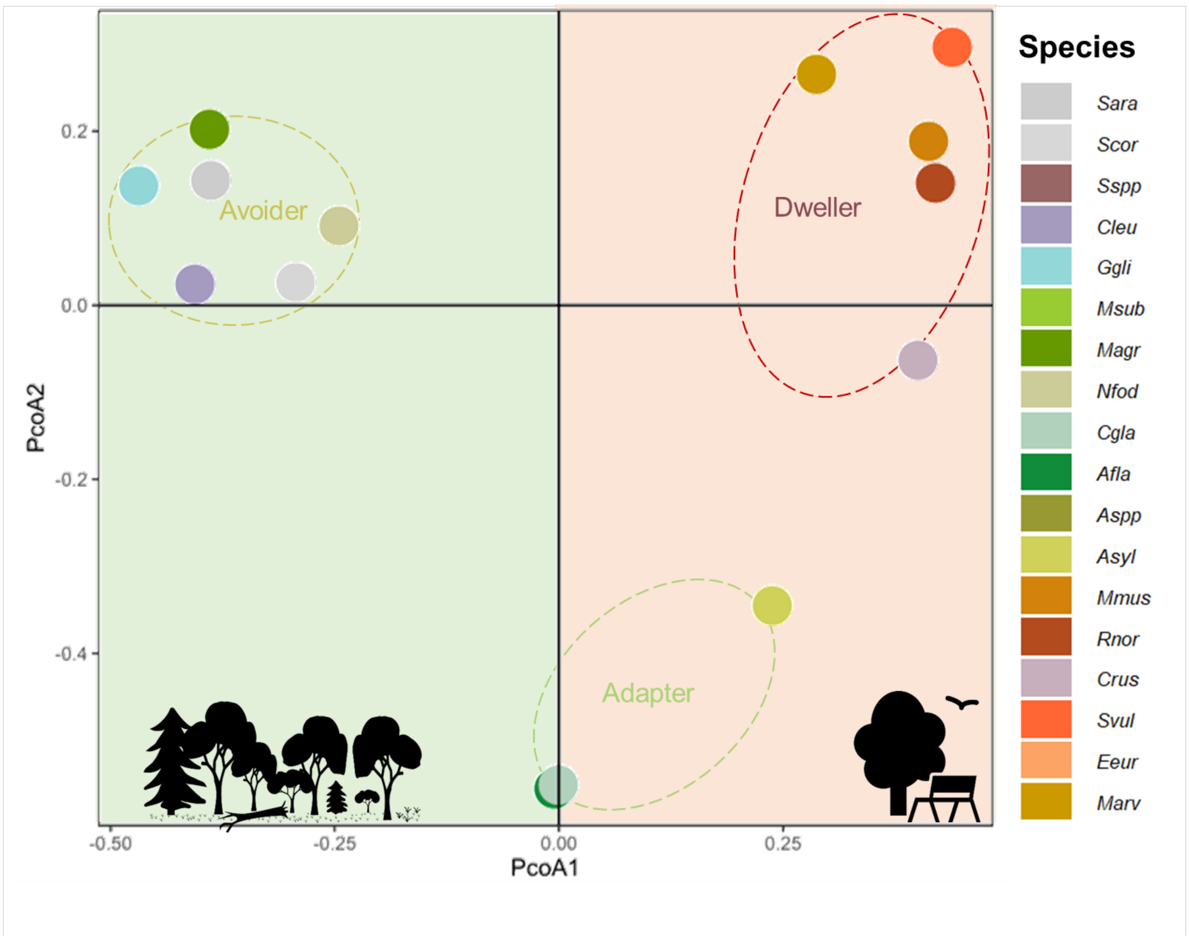


**Supplementary Fig. 2. Principal Coordinates Analysis (PCoA) based on the distribution of small mammal species across study sites,** using both abundance and presence/absence data (Bray-Curtis dissimilarities), performed with the vegan package in R (Oksanen et al. 2020). This analysis highlights that species occurring across both the anthropization gradient tend to cluster separately from species restricted to a single habitat type. Background colors reflect habitat types: green indicates non-urban sites, and red corresponds to urban sites. icons/logos and their positions along the first PCoA axis illustrate a gradient of anthropization level, reflecting their habitat affiliation and tolerance to anthropogenic environments. This gradient allowed us to define three ecological species types based on their urban adaptation: avoiders (avoiding urban environments), adapters (found across all habitat types), and dwellers (restricted to urban parks). Species codes: *Asyl = Apodemus sylvaticus, Afla = Apodemus flavicollis, Cgla = Clethrionomys (Myodes) glareolus,Crus = Crocidura russula, Cleu = Crocidura leucodon, Mmus = Mus musculus, Rnor = Rattus norvegicus, Ggli = Glis glis, Msub = Microtus subterraneus, Marv = Microtus arvalis, Magr = Microtus agrestis,, Nfod = Neomys fodiens, Svul = Sciurus vulgaris, Eeur = Erinaceus europaeus,, Sara = Sorex araneus, Scor = Sorex coronatus.*

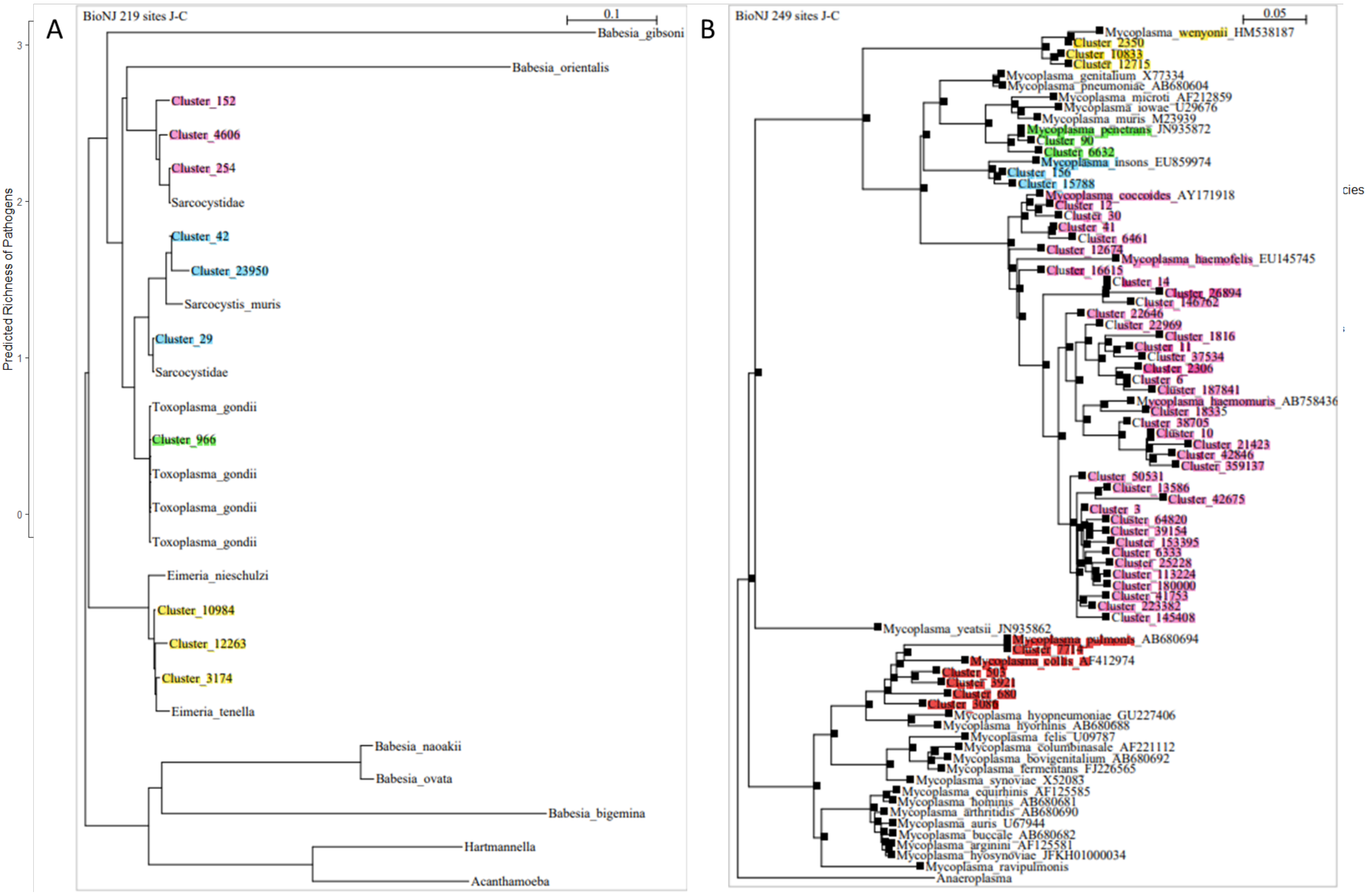


Supplementary Fig. 3. Phylogenetic Trees of *Sarcocystidae* and *Mycoplasma* Based on 16S V4 Sequences. FASTA sequences obtained from sequencing files were used to construct phylogenetic trees. OTUs detected by FROGS (Escudié et al. 2018) were assigned to reference databases via Silva. Some unknown OTUs, non-bacterial or unresolved at the species level, were subjected to BLAST analysis to confirm their affiliation with *Apicomplexa* or *Mycoplasma*. Closely related sequences were used as references. Some sequences affiliated with Sarcocystidae did not show significant matches, suggesting the presence of potentially undescribed strains. Alignments were performed using SeaView (Gouy, Guindon, et Gascuel 2010) and MUSCLE, and the phylogenetic tree was constructed using the BioNJ method and the Jukes-Cantor distance model. **A.** ***Sarcocystidae***: The **g**reen cluster corresponds to *Toxoplasma gondii*, and the yellow clusters are identified as *Eimeria*. The blue cluster groups sequences affiliated with *Sarcocystis G. spp.*, while the pink cluster represents another group within Sarcocystidae. **B.** *Mycoplasma*: Two unknown OTUs were assigned to *Mycoplasma penetrans* (zoonotic) in the green cluster, while the other sequences were closely related to *Mycoplasma coccoides* and *Mycoplasma haemomuris*, which are pathogenic to rodents but not zoonotic.

**Supplementary Figure 4. Boxplots of predicted individual pathogen richness from zero-inflated Poisson GLMs, according to: A. Sites, ordered along an anthropization gradient ;B. periods, C. small mammal host species.** Each point represents an individual; overlapping points may be hidden due to identical richness values. Urban park forests are coded as follows: FRPLTO (red) – Lyon, Parc de la Tête d'Or (Rhône); FRPDLL (orange) – Marcy l'Étoile, Domaine Lacroix-Laval (Rhône); rural managed or protected forests are FRFCOR – Cormaranche-en-Bugey (Ain); FRFGRI – Arvière, La Griffe au Diable (Ain); FRFMIG – Mignovillard (Jura); FRFGLA – Esserval-Tartre, La Glacière (Jura), shown in bluish-green tones; seasons are coded as s = spring (grey) and f = fall (green), followed by the sampling year and ordered chronologically; host species are color-coded according to behavior: urban dwellers in red hues, adapters and avoiders in green tones, shrews in purple; species codes are as follows: Asyl = Apodemus sylvaticus; Afla = Apodemus flavicollis; Cgla = Clethrionomys glareolus; Crus = Crocidura russula; Cleu = Crocidura leucodon; Mmus = Mus musculus; Rnor = Rattus norvegicus; Ggli = Glis glis; Marv = Microtus arvalis.

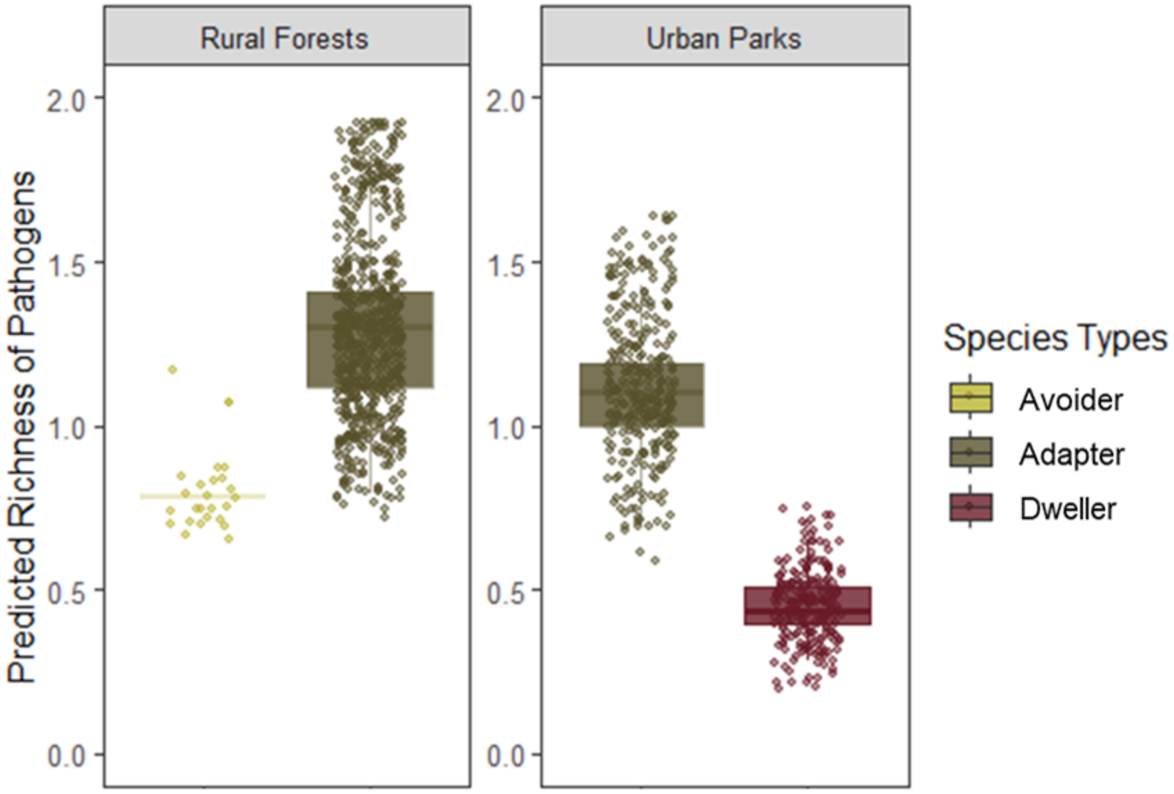


**Supplementary Figure 5. Boxplots of predicted individual pathogen richness from zero-inflated Poisson GLMs, according to the ecological types of small mammals along the anthropization gradient:** *avoiders* (yellow), which avoid urban areas; *adapters* (green), present across all habitats; and *dwellers* (red), found exclusively in urban parks. Comparisons were made between two habitat categories: rural forests (including both protected and managed forests, grouped due to the low number of *avoider* individuals in each) and urban parks. Each point represents an individual; overlapping points may be hidden due to identical richness values.

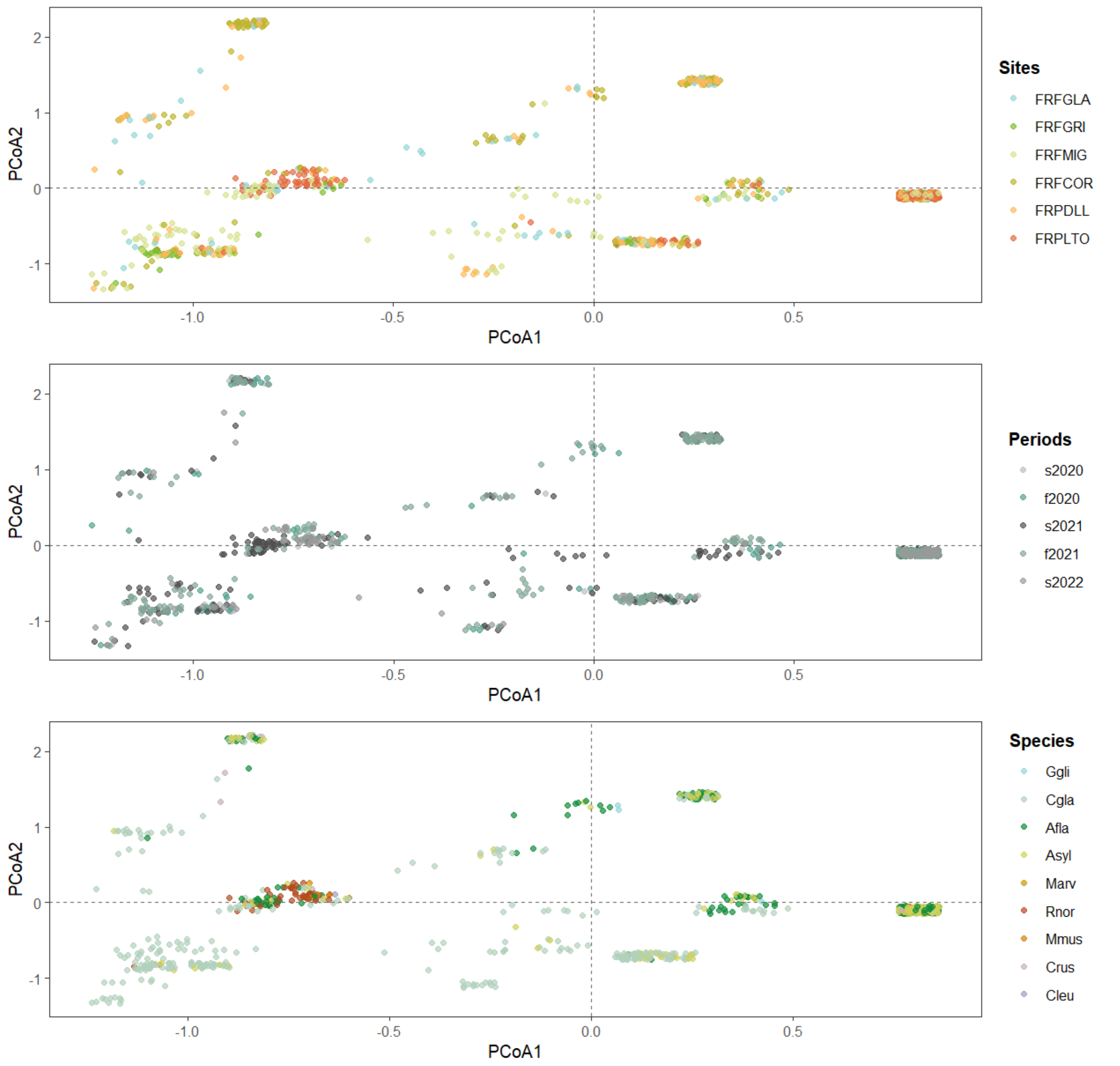


**Supplementary Figure 6. Pathogen community composition (presence of at least one pathogen among the 16 tested) based on Jaccard dissimilarities visualized by PCoA, colored according to site, sampling period, and host species. PCoA axis 1 and axis 2 explain 35.2% and 14.7% of the variance, respectively. Each point represents an individual.** Urban park forests are coded as: FRPLTO (red) – Lyon, Parc de la Tête d'Or (Rhône); FRPDLL (orange) – Marcy l'Étoile, Domaine Lacroix-Laval (Rhône); rural managed or protected forests are: FRFCOR – Cormaranche-en-Bugey (Ain), FRFGRI – Arvière, La Griffe au Diable (Ain), FRFMIG – Mignovillard (Jura), and FRFGLA – Esserval-Tartre, La Glacière (Jura), shown in bluish-green tones. Sampling periods are coded as *s* = spring (grey) and *f* = fall (green), followed by the year and ordered chronologically. Host species are color-coded by ecological behavior: *dwellers* in red hues, *adapters* and *avoiders* in green tones, and shrews in purple. Species codes: *Asyl* = *Apodemus sylvaticus*; *Afla* = *Apodemus flavicollis*; *Cgla* = *Clethrionomys glareolus*; *Crus* = *Crocidura russula*; *Cleu* = *Crocidura leucodon*; *Mmus* = *Mus musculus*; *Rnor* = *Rattus norvegicus*; *Ggli* = *Glis glis*; *Marv* = *Microtus arvalis*. Adonis2 results from the vegan package revealed significant effects of host species ($R^2$ = 17.4%, $p < 0.001$), site ($R^2$ = 6.9%, $p < 0.001$), period ($R^2$ = 4.4%, $p < 0.001$), and age class ($R^2$ = 0.4%, $p = 0.002$), but not sex. Interactions between species and sites were also significant ($R^2$ = 2.3%, $p < 0.001$). Post-hoc comparisons confirmed that most levels within each factor differed significantly (see sup table 3). These findings indicate that pathogen communities are highly structured by environmental conditions and host identity, with some individual variation (e.g., age), though the variation was too continuous to define distinct clusters (ellipses not shown).

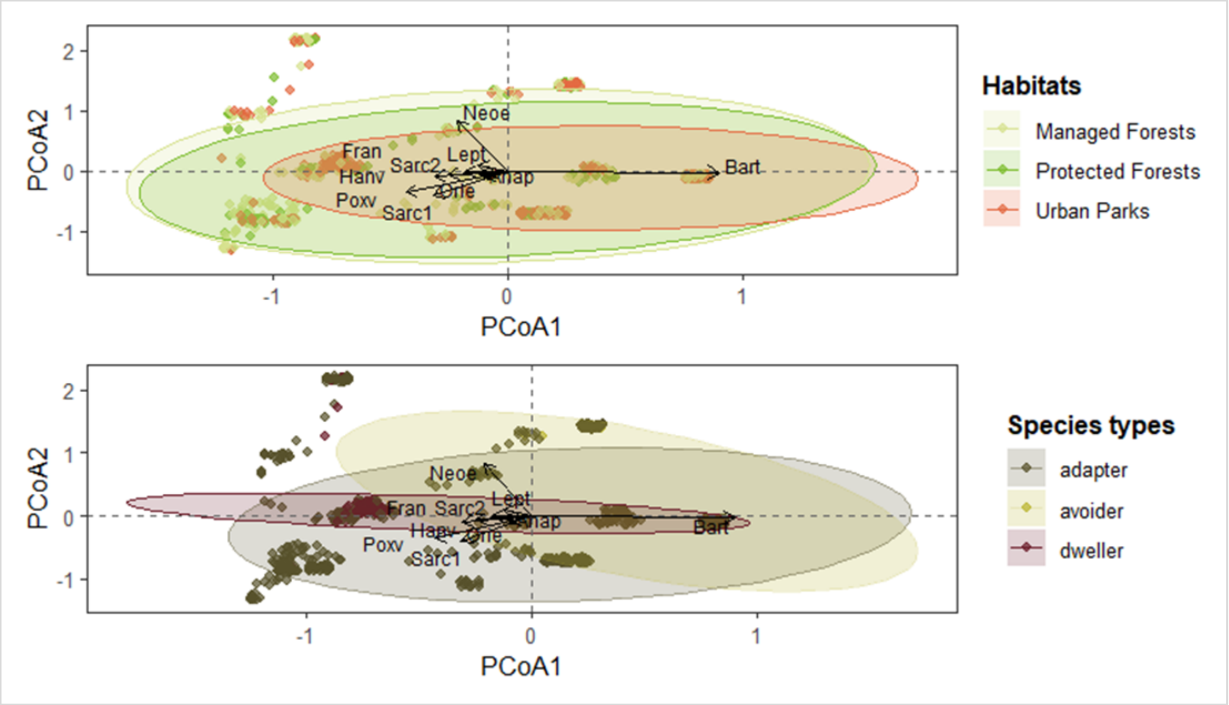


**Supplementary Figure 7. Pathogen community composition (presence of at least one of the 16 pathogens) based on Jaccard dissimilarities, visualized using PCoA , with axis 1 and axis 2 explain 35.2% and 14.7% of the variance, respectively**. Each point represents an individual, and ellipses indicate 95% confidence intervals around the centroids of the different modalities. Points and ellipses are colored according to habitat type along a management gradient—protected forests (green), managed forests (yellow), and urban parks (red)—and host ecological types based on distribution: avoiders (light green; avoiding urban areas), adapters (dark green; present across all habitats), and dwellers (red; restricted to urban parks). PERMANOVA results (see Supplementary Table 4) confirmed significant differences in pathogen community composition across both species ecological types ($R^2 = 0.034$, $p < 0.001$) and habitats ($R^2 = 0.035$, $p < 0.001$).

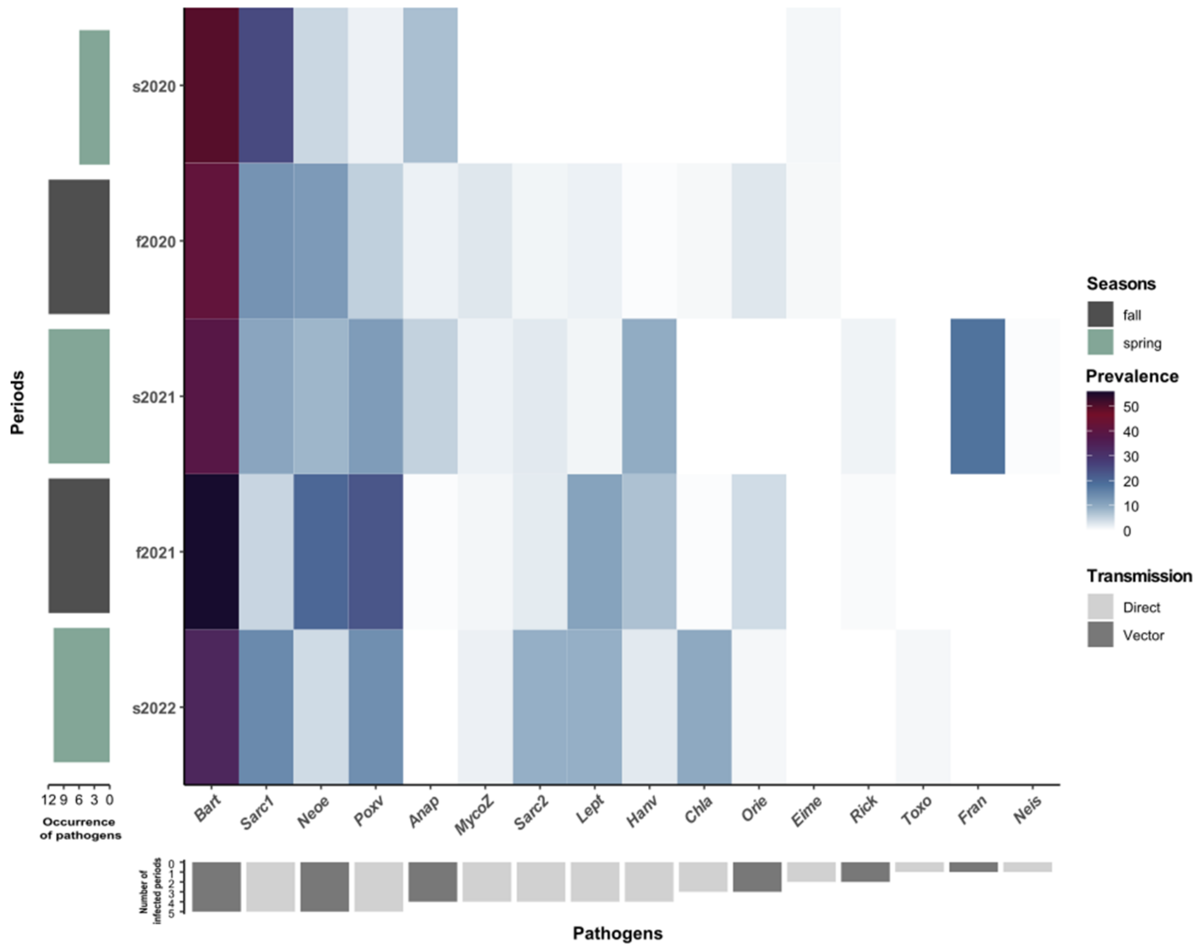


Supplementary Figure 8. Heatmap of pathogen (sero)prevalence across sampling periods. Sampling periods are ordered chronologically; *s* = spring (light green) and *f* = fall (dark green). Color gradients from light blue to dark purple indicate increasing (sero)prevalence. Vertical bar charts on the left represent the number of pathogens detected per period. Inverted horizontal bar charts below the heatmap indicate the number of periods in which each pathogen was detected. Transmission modes are indicated by bar colors: dark grey for vector-borne transmission, and light grey for direct transmission (via contact and/or environmental exposure).
